# The astrocytic secretome increases adult hippocampal neurogenesis and spatial memory in health and disease

**DOI:** 10.64898/2026.09.26.754656

**Authors:** Marta Vilademunt, Charline Carron, Frederic Cassé, Kevin Richetin, Elias Gebara, Kyllian Ginggen, Jonathan Nicolet-dit-Felix, Luana Piano-Vieira, Nicolas Toni

## Abstract

**INTRODUCTION:** Adult hippocampal neurogenesis (AHN) plays an important role in memory and is regulated by astrocytes. Since astrocytic function deteriorates with pathological ageing and Alzheimer’s disease (AD), we hypothesize that the astrocytic secretome may rescue AHN and cognition.

**METHODS:** We tested the effect of astrocyte-conditioned solution (ACS) on AHN and memory performances in young and aged mice and in two models of AD.

**RESULTS:** ACS increased AHN and improved performance in the object location test in young and aged wild-type mice. In THY-Tau22 mice, a model of tau pathology, ACS restored AHN and cognitive performance. In APP/PS1 mice, a model of amyloid pathology, ACS reduced gliogenesis, and rescued deficits in AHN and spatial memory. In addition, ACS attenuated circulating pro-inflammatory cytokines and the reaction of plaque-associated microglia.

**DISCUSSION:** The astrocytic secretome is a promising, niche-targeted strategy to enhance hippocampal plasticity and memory across ageing and AD.

## 1. Background

Cognitive decline is a hallmark of ageing and neurodegeneration and closely reflects the progressive loss of hippocampal plasticity^1^. One prominent form of such plasticity is adult hippocampal neurogenesis (AHN), which results in the lifelong generation of new neurons in the dentate gyrus^2^. AHN plays a role in learning, memory, and emotional regulation^3–8^ that is mediated in particular by immature neurons that undergo a critical period at around 4-6 weeks after cell division, during which they display increased plasticity^6, 9^. AHN declines sharply with age^10–12^, which is caused mainly by a reduction in neural stem/progenitor cell density and self-renewing capacity and is accompanied by a decrease in spatial memory performances^1, 13–19^. AHN and memory performances are further disrupted in neurodegenerative conditions, underscoring the need to identify strategies that preserve or restore neurogenesis^20^.

Interventions to restore AHN in aged mice are often accompanied by improvements in learning and memory^21–25^. Similarly, the experimental enhancement of AHN in AD models ameliorates cognitive performances^26–32^.

Interestingly, the two main pathological hallmarks of AD—amyloid-β plaques and tau pathology—have each been linked to impaired neurogenesis^33–36^, alongside microglial activation and chronic neuroinflammation^37^. This inflammatory response further disrupts the neurogenic niche, thereby exacerbating AHN impairment. Astrocytes are important components of the neurogenic niche^38^. That regulate AHN by releasing secreted factors^39, 40^, providing structural support of newborn neurons^41, 42^, regulating the local microenvironment required for synaptic integration^43^ and reducing inflammation^44^. However, the astrocytic structure and function is itself profoundly altered by ageing and neurodegenerative disease: Ageing is associated with a loss of astrocytic homeostasis, including structural atrophy, functional and metabolic impairments, and the upregulation of transcriptomic signatures linked to neurotoxicity^45–47^. In AD, astrocytes undergo additional pathological changes, with transcriptomic analyses revealing downregulation of homeostatic genes and dysregulation of key pathways^48, 49^. Although the consequences of these alterations for the neurogenic niche remain incompletely understood, the experimental expression of a pathological form of tau in astrocytes in the hilus of the dentate gyrus is sufficient to impair AHN and memory performances^50^. Thus, the astrocytic support to AHN plays an important role in hippocampal function and is compromised in AD.

Most approaches to enhance AHN in ageing, and neurodegenerative contexts have focused on directly stimulating NSCs, targeting disease-specific mechanisms or experience-dependent interventions such as physical exercise^3, 29^ and environmental enrichment^51^. However, strategies aimed at compensating for functional deficits of the neurogenic niche itself remain largely unexplored. Given the central role of astrocytes in coordinating niche homeostasis and neurogenesis, the astrocytic secretome represents a promising and under investigated therapeutic avenue.

In the present study, we investigated the effects of the astrocytic secretome in the form of astrocyte-conditioned solution on AHN, neuroinflammation, and cognitive performance under physiological conditions and in two complementary mouse models of AD: the THY-Tau22 model of tauopathy^52^ and the APPswe/PS1de9 (APP/PS1) model of amyloidosis.

## 2. Material and Methods

### Study Design and Data Handling

Animals were randomly assigned to experimental groups, ensuring an equal distribution of males and females within each litter and randomization across litters to minimize litter-specific effects. To reduce cage effects, randomization was performed so that each cage contained an equal proportion of animals from each treatment group. Data collection and processing were conducted in a randomized order, and experimenters were blinded to group assignments during data analysis. Sample sizes for all groups were determined using an a priori power analysis to ensure sufficient statistical power.

### Animals

All experimental procedures were approved by the Cantonal Veterinary Authorities (Vaud, Switzerland) and conducted in accordance with the European Communities Council Directive of 24 November 1986 (86/609/EEC). Healthy C57BL/6J mice were used, with adult males aged 8 weeks at the start of the experiments. For experiments shown in Figure 2, 18-month-old C57BL/6J mice were used. C57BL/6J mice were obtained from Janvier (Le Genest Saint Isle, France). THY-Tau22 mice, kindly provided by Claire Rampon (Research Center on Animal Cognition, Toulouse) were bred in-house. Mice aged 11–14 months were used. APP/PS1 mice, kindly provided by Prof. Joannes Graef (EPFL, Lausanne), were bred in-house; females aged 4 months ± 1 week and males aged 6 months ± 1 week were used, as they exhibit comparable amyloid loads. Postnatal day 1–2 C57BL/6J pups were used for primary cultures. All animals were maintained under a 12-hour light/dark cycle with ad libitum access to food and water, in a controlled environment at 23°C ± 1°C.

### Cell culture

#### Mouse astrocyte primary cultures

Astrocytes were prepared from postnatal day 1–2 WT mice (both female and male pups), as previously described (Sultan et al., 2015). Briefly, pups were decapitated, brains were rapidly collected, and meninges were carefully removed. Hippocampi and cortices were dissected, mechanically triturated, and seeded onto 25 cm² flasks in DMEM with GlutaMAX™ (Gibco/Life Technologies, 10569-010), supplemented with 10% horse serum (Gibco/Life Technologies, 16050-122) and 1% PSF (Gibco/Life Technologies 15240062). Cultures were maintained for one week in a humidified 5% CO₂ incubator at 37°C. Flasks were shaken every 2 days to separate microglia from astrocytes and washed with HBSS (Gibco/Life Technologies, 14025-092).

### In vivo experiments

#### DNA extraction and Genotyping

Genotyping was required for the APP/PS1 mouse line, as this line is maintained in a heterozygous state. Biopsy samples were digested overnight at 55°C with shaking (300 rpm) in lysis buffer containing proteinase K. The following day, DNA was extracted by centrifugation (13,000 rpm, 10 min, room temperature), followed by precipitation with isopropanol, washing with 70% ethanol, air-drying, and resuspension in sterile water. PCR amplification of APP and PS1 genes was performed using 2 μL of DNA template in a 25 μL reaction containing Taq polymerase mix (5X GREEN GoTaq Flexi Buffer, Promega), primers for each target gene, and water. The cycling program consisted of an initial denaturation at 94°C for 3 min, followed by 34 cycles of 94°C for 30 s, 60°C for 1 min, and 72°C for 1 min, with a final extension at 72°C for 2 min. For genotyping, 15 μL of each PCR product was resolved on a 1% agarose gel containing SYBR Safe DNA stain in 1× TAE buffer. Gels were cast with wells, allowed to solidify, and electrophoresed for ∼40 min. DNA bands were visualized under UV light to determine the presence of the target genes.

#### Astrocyte-conditioned Solution (ACS)

Mouse primary astrocyte cultures were maintained in flasks for at least 10 days to achieve confluency. Prior to experiments, cultures were washed twice with HBSS (Gibco/Life Technologies 14025-092) to remove residuals of cell culture medium and incubated in 2 mL of Tyrode solution per flask (composition: HEPES 10 mM, NaCl 145 mM, KCl 5.4 mM, CaCl₂ 1.8 mM, MgCl₂ 0.8 mM, Glucose 10 mM, q.s. ddH₂O) at 37°C for 24 hours. The conditioned solution was then collected and filtered through a 0.22 µm filter (Millipore) to remove cellular debris.

#### Injections

Mice received intraperitoneal injections of 200 μL of ACS, pre-warmed to 37°C, prepared from a different astrocyte culture each day. BrdU was dissolved in Tyrode solution and injected intraperitoneally at 100 mg/kg (Sigma-Aldrich, Buchs, Switzerland) three times on the day of injection, at 2-hour intervals.

#### Brain tissue and serum collection

Mice received a lethal intraperitoneal injection of pentobarbital (10 mL/kg; Sigma-Aldrich, Buchs, Switzerland). For APP/PS1 serum samples: Once anaesthesia was confirmed, the thoracic area was disinfected with 70% ethanol. A maximum of 0.4 mL blood was then collected via intracardiac puncture using a 23–27G needle and 1 mL syringe. Samples were allowed to clot upright at room temperature for 10 minutes before being centrifuged at 3,500 rpm (∼1,900 × g) for 10 minutes. Up to 90 µL of clear serum was carefully collected, flash-frozen on dry ice, and initially stored at −20 °C; samples were then transferred to −80 °C for long-term storage. For histology, animals were perfused with 0.9% NaCl for 3 minutes, followed by 4% paraformaldehyde (PFA) for 3 minutes (Sigma-Aldrich, USA) dissolved in 0.1 M Phosphate Saline Buffer (PBS, pH 7.4). Their brains were dissected, fixed overnight at 4°C in 4% PFA, and then incubated for 24 hours in a 30% sucrose solution at 4°C (Sigma-Aldrich, USA). Then brains were transferred in a 30% sucrose solution for 3 days before being frozen at −20 °C until slicing. 35 μm (Figure 4-7) or 40 μm (Figures 1-3) thick coronal sections were prepared using a cryostat (Leica MC 3050S) and were preserved in cryoprotectant (30% ethylene glycol + 25% glycerin in 0.1 M PBS) at −20°C until immunofluorescence staining.

**Figure 1.**
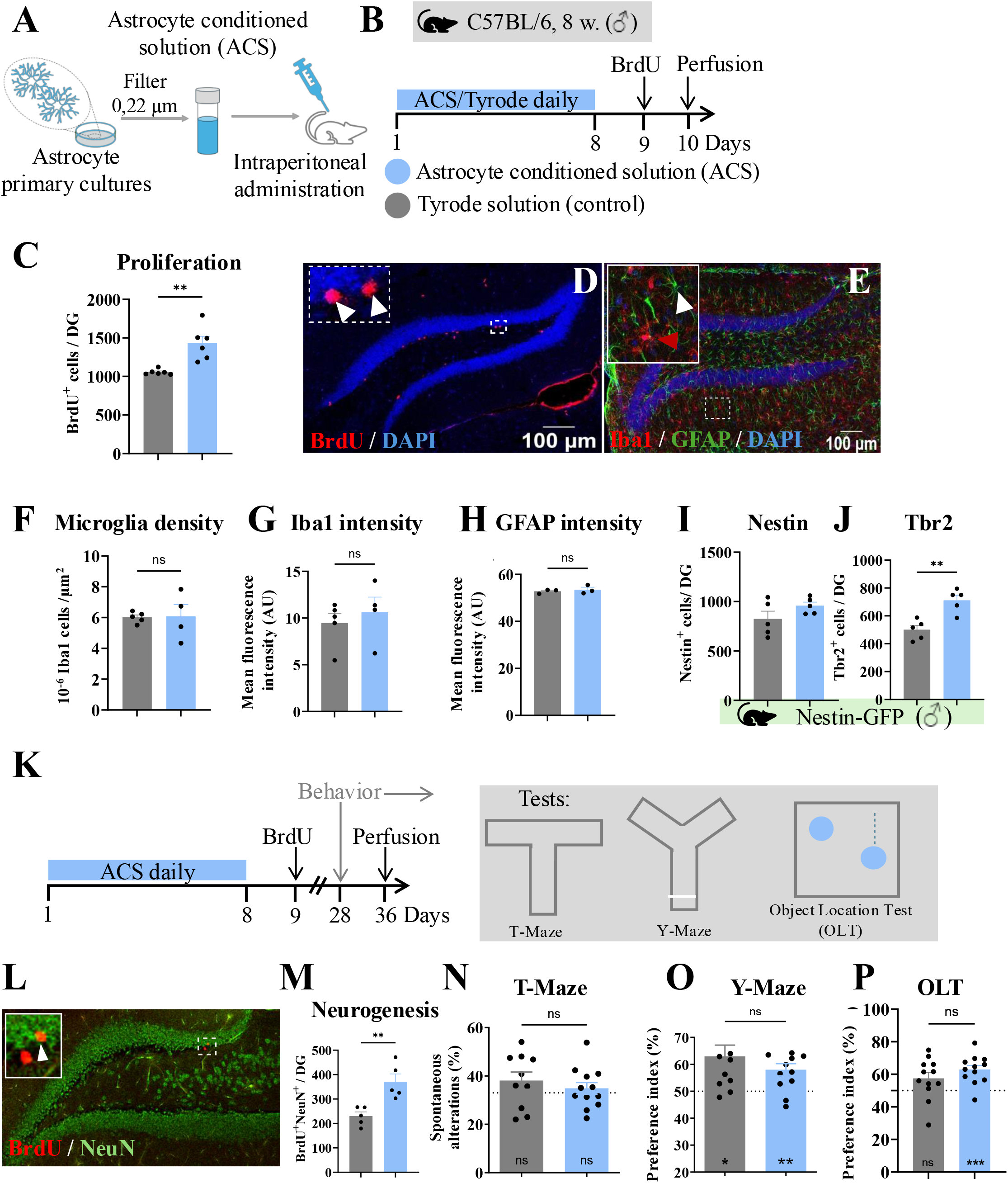
ACS enhances neurogenesis and improves memory in healthy conditions. **A)** Schematic representation of the obtention of astrocyte conditioned solution (ACS) **B)** Experimental timeline. **C)** Quantification of BrdU^+^ proliferative cells in the total dentate gyrus (n = 12, t = 4.15, p = 0.002). **D-E)** Representative confocal microscopy images showing BrdU (Red) and DAPI (blue) (**D**) or Iba1 (red), GFAP (green) and DAPI (blue) (**E**). Insets show higher magnification of labelled cells with white arrows indicating examples of marker-positive cells. **F)** Quantification of Iba1^+^ microglia density (n = 9, t = 0.098, p = 0.924). **G-H)** Iba1^+^ microglia (**G:** n = 9, t = 0.623, p = 0.553) and GFAP^+^ astrocyte (**H:** n = 6, t = 0.601, p = 0.58) mean fluorescence intensity. **I)** Number of Nestin^+^ cells in the dentate gyrus of Nestin-GFP mice (n = 10, t = 1.525, p = 0.66). **J)** Tbr2^+^ cells in the dentate gyrus of Nestin-GFP mice (n = 10, t = 4.324, p = 0.0025). **K)** Experimental design. **L)** Confocal microscopy image of the dentate gyrus, BrdU (red) and NeuN (green). **M)** Quantification of BrdU⁺/NeuN⁺ double-labelled cells in the total dentate gyrus (n = 12, t = 3.908, p = 0.0045). **N)** Quantification of the spontaneous alteration on the T-Maze (n = 24, t = 0.754, p = 0.46). One sample t-test (wt: t = 1.415, p = 0.19, ACS: t = 0.77, p = 0.46) **O)** Preference index in the Y-Maze (n = 24, t = 1.055, p = 0.3). One sample t-test (wt: t = 3.063, p = 0.01, ACS: t = 3.535, p = 0.0047) **P)** Preference Index in the Object Location Test (n = 24, t = 1.24, p = 0.23). One sample t-test (wt: t =2.042, p = 0.0659, ACS: t = 5.075, p = 0.0004). Data is shown as mean ± SEM. Statistics: unpaired t-test. *p ≤ 0.05, **p ≤ 0.01, ****p ≤ 0.0001. Scale bar = 100 μm.

**Figure 2.**
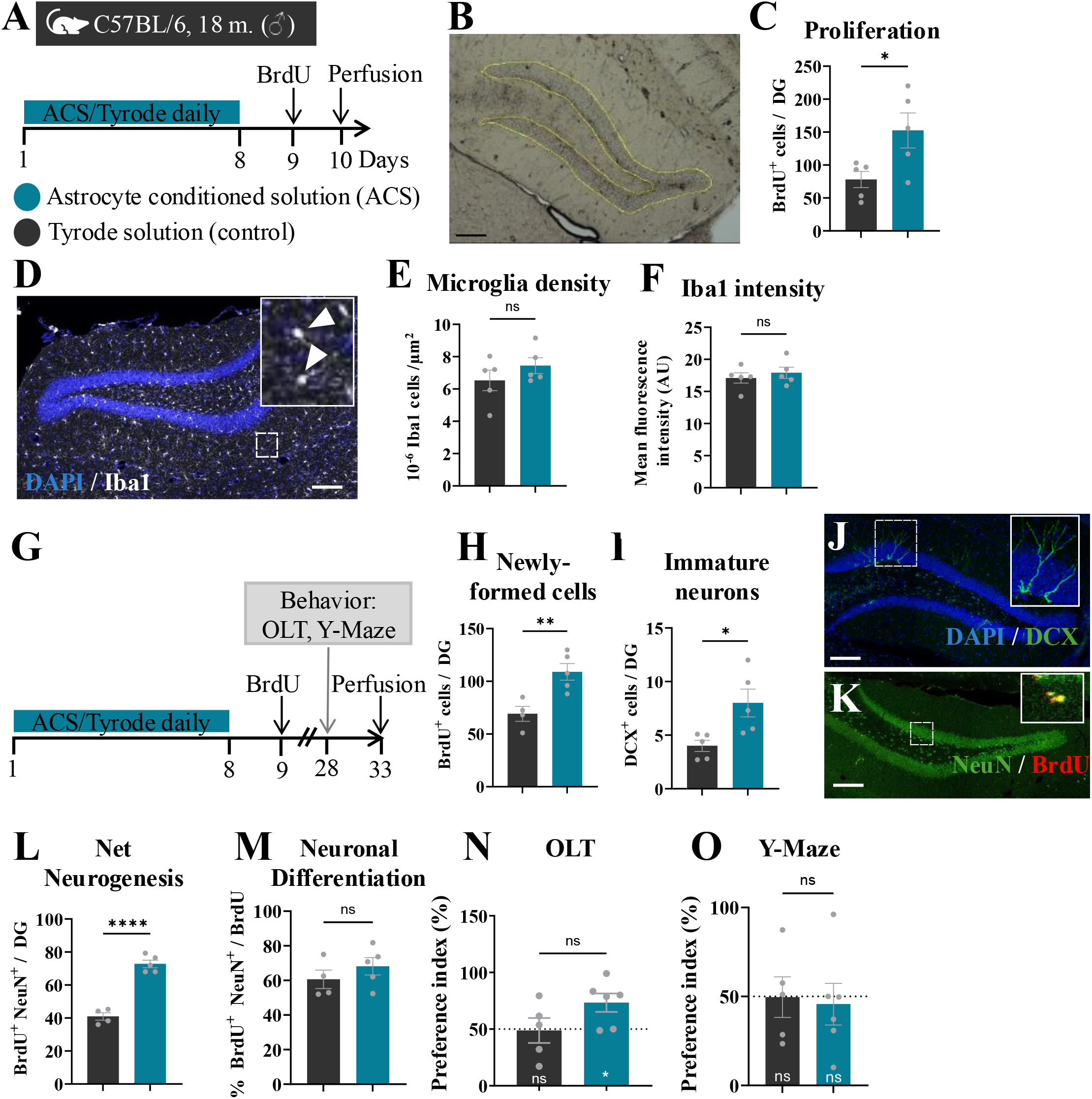
ACS enhances neurogenesis and memory performance in aged mice. **A)** Experimental timeline. **B)** Microscopy image of mouse dentate gyrus. **C)** Quantification of BrdU^+^ cells in the dentate gyrus (n = 10, t = 2.55, p = 0.03) **D)** Representative confocal microscopy image. DAPI (blue), Iba1 (white). Insets show higher magnification of labelled cells with white arrows indicating examples of marker-positive cells. **E)** Iba1^+^ cells in the dentate gyrus (n = 10, t = 1.133, p = 0.290). **F)** Iba1^+^ mean fluorescence intensity (n = 10, t = 0.672, p = 0.520). **G)** Experimental timeline. **H)** BrdU^+^ cells in the subgranular zone of the dentate gyrus (n = 10, t = 3.66, p = 0.008). **I)** DCX^+^ cells in the dentate gyrus (n = 10, t = 2.857, p = 0.021). **J, K)** Representative confocal images. DAPI (blue), DCX(green), NeuN (green), BrdU (red). **L)** Quantification of BrdU⁺/NeuN⁺ double-labelled cells in the dentate gyrus (n = 10, t = 9.781, p < 0.0001). **M)** Percentage of BrdU labelled cells that are also labelled with NeuN^+^ (n = 10, t = 1.019, p = 0.342). **N)** Preference index in the Objection Location Test (n = 10, t = 1.833, p = 0.100). **O)** Preference index in the Y-Maze (n = 11, t = 0.238, p = 0.82). Data is shown as mean ± SEM. Statistics: unpaired t-test. *p ≤ 0.05, **p ≤ 0.01. Scale bar = 100 μm.

**Figure 3.**
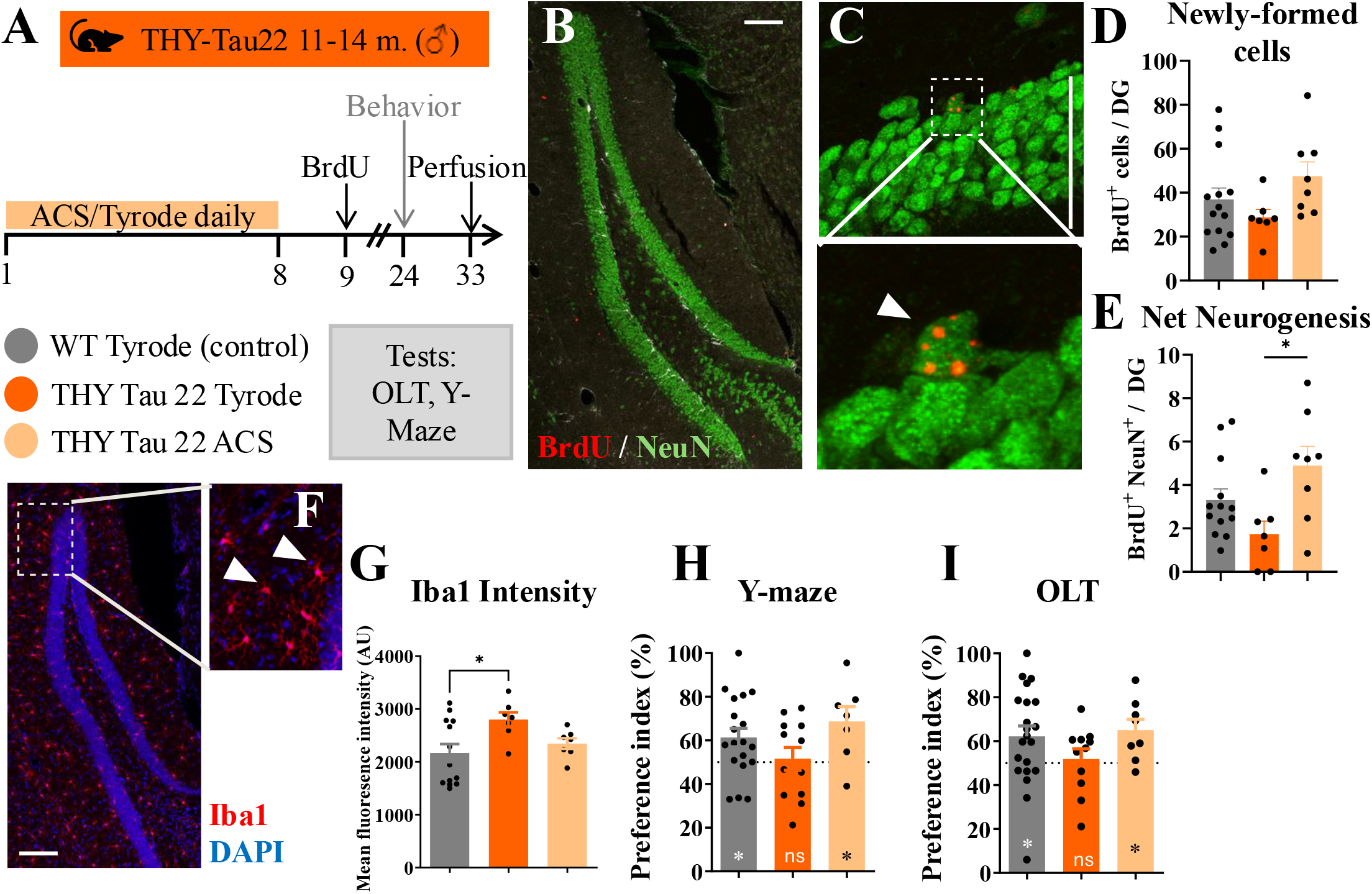
Astrocyte conditioned solution rescues neurogenic and cognitive loss in a mouse model of tau pathology. **A)** Experimental timeline. **B-C)** Representative confocal images. BrdU (red), NeuN (green). Insets show higher magnification of labelled cells with white arrows indicating examples of marker-positive cells. **D)** BrdU^+^ cells in the subgranular zone of the dentate gyrus (n = 30, F(2,26) = 2.165, p = 0.135). **E)** Quantification of BrdU⁺/NeuN⁺ double-labelled cells in the dentate gyrus (n = 29, F(2, 25) = 4.603, p = 0.02). **F)** Representative confocal image. Iba1 (red), DAPI (Blue). Inset show higher magnification of labelled cells with white arrows indicating examples of Iba1-immunostained cells. **G)** Iba1 mean fluorescence intensity (n = 28, F(2,24) = 3.768, p = 0.038). **H)** Preference index of the closed arm in Y-Maze (n = 38, F(2,35) = 2.107, p = 0.14). **I)** Preference index in the Object Location Test (n = 40, F(2,37) = 1.424, p = 0.33). Data is shown as mean ± SEM. Statistics: One-way ANOVA with Tukey’s multiple comparisons test. *p ≤ 0.05, **p ≤ 0.01, ****p ≤ 0.0001. Scale bar = 100 μm.

**Figure 4.**
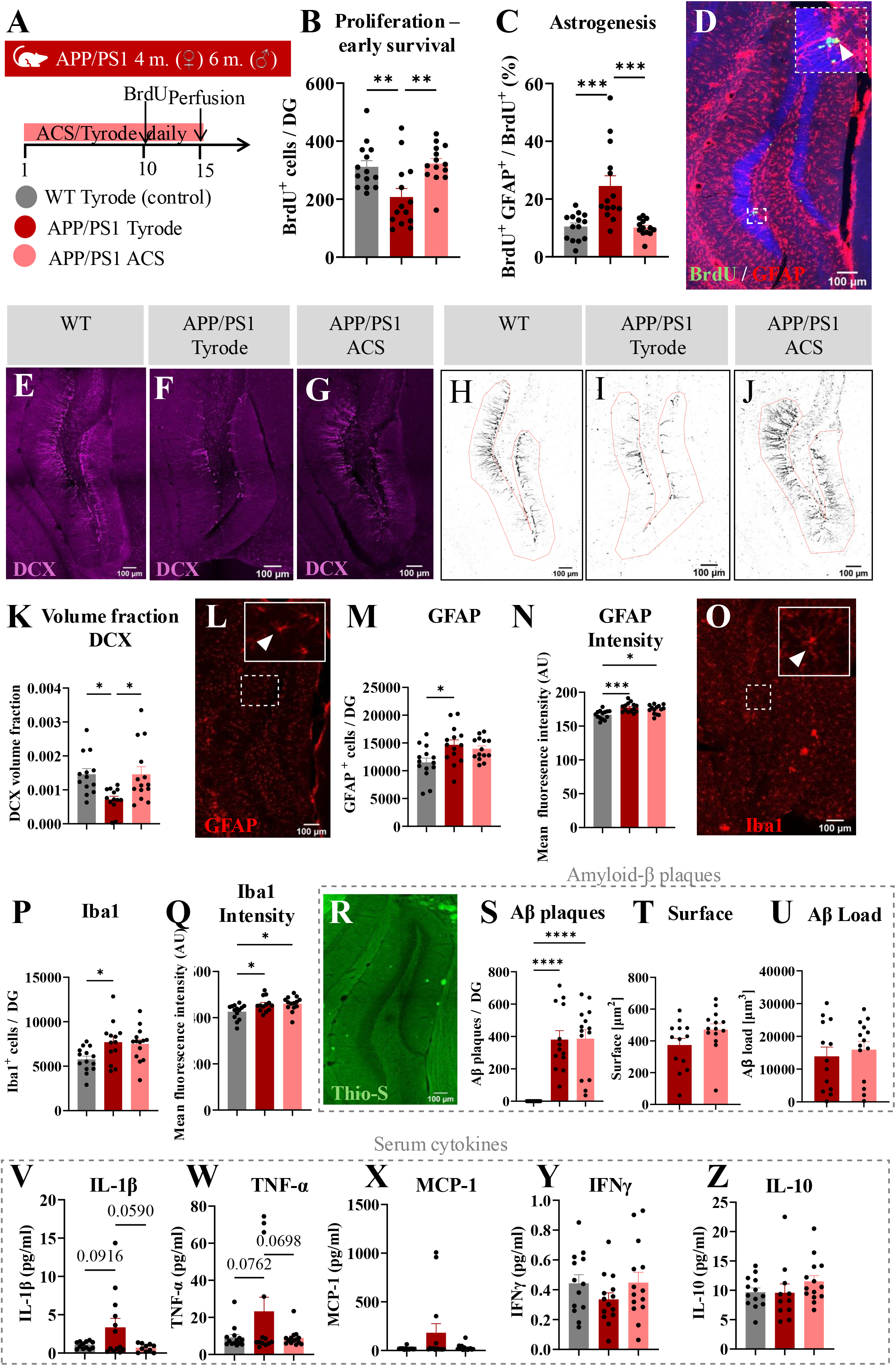
ACS rescues neurogenesis and reduces neuroinflammation in a mouse model of amyloidosis. **A)** Experimental timeline. **B)** BrdU^+^ cells in the subgranular zone of the dentate gyrus (n =42, F(2,39) = 7.46, p = 0.0018). **C)** Percentage of double-labelled cells with BrdU^+^ and GFAP^+^ from the total BrdU population, quantified in the subgranular zone of the dentate gyrus (n = 41, F(2,38) = 13.12, p <0,0001). **D)** Representative confocal image. BrdU (green), GFAP (red). Inset show higher magnification of labelled cells with white arrow indicating an example of a BrdU-positive cell. **E-J)** Representative confocal images showing DCX⁺ immunolabeling (purple) and the corresponding threshold-processed images used for DCX⁺ volume fraction quantification (black) within the analyzed region of interest (ROI, outlined in red). **K)** Proportion of the dentate gyrus occupied by DCX⁺ immunolabeling defined as DCX volume fraction (n = 40, F(2,37) = 5.870, p = 0.0061). **L)** Representative confocal image. GFAP (red). Inset show higher magnification of labelled cells with white arrow indicating an example of a GFAP-positive cell. **M)** Quantification of GFAP^+^ cells in the dentate gyrus (n = 42, F(3,39) = 4.8, p = 0.01). **N)** Mean fluorescence intensity of GFAP immunolabeling (n = 42, F(2,39) = 9.57, p = 0.0004). **O)** Representative confocal image. Iba1 (red). Inset show higher magnification of labelled cells with white arrow indicating an example of a Iba1-positive cell. **P)** Quantification of Iba1^+^ cells in the dentate gyrus (n = 41, F(2,38) = 4.32, p = 0.02). **Q)** Mean fluorescence intensity of Iba1 immunolabelling (n = 41, F(2,38) = 4.96, p = 0.01). **R)** Representative confocal image. Thioflavin-S (Thio-S, green). **S)** Quantification of amyloid-β plaques in the dentate gyrus (n = 40, F(2,37) = 22.87, p < 0.0001). **T)** Quantification of amyloid-β plaques’ surface [µm^2^] (n = 27, t = 1.75, p = 0.09). **U)** Amyloid-β load in the dentate gyrus (n = 27, t = 0.54, p = 0.59)**. V)** Levels of IL-1β in serum (n =35, F(2,32) = 3.58, p = 0.039). **W)** Levels of TNF-α in serum (n = 41, F(2,38) = 3.38, p = 0.044). **X)** Levels of MCP-1 in serum (n = 42, F(2,39) = 2.8, p = 0.07). **Y)** Levels of IFNγ in serum (n = 41, F(2,38) = 1.22, p = 0.30). **Z)** Levels of IL-10 in serum (n = 38, F(2,35) = 1.09, p = 0.35). Data is shown as mean ± SEM. Statistics: One-way ANOVA with Tukey’s multiple comparisons test or unpaired T-test. *p ≤ 0.05, **p ≤ 0.01, ****p ≤ 0.0001. Scale bar = 100 μm.

#### Immunohistochemistry

One out of every six coronal sections containing hippocampal tissue was selected systematically to cover the entire rostro-caudal extent of the dentate gyrus for immunostaining. For detection of BrdU-positive cells and co-labeling with either NeuN or GFAP, sections underwent formamide-based pretreatment (50% formamide, 10% 20X SSC, 40% MilliQ water) at 60°C for 2 hours, followed by DNA denaturation in 2 M HCl for 30 minutes at 37°C. Sections were then rinsed in 0.1 M borate buffer (pH 8.5) for 15 minutes and washed six times in 0.1 M PBS for 10 minutes each. Sections were incubated in blocking solution containing 0.3% Triton X-100 and 5% normal donkey or goat serum in PBS for 1 hour at room temperature to prevent nonspecific binding. Subsequently, sections were incubated under gentle agitation at 4°C for 48 hours with mouse anti-BrdU (1:300, Abcam ab6326) or rat anti-Brdu (1:500, Abcam ab6326) and rabbit anti-NeuN (1:1000, Abcam EPR12763) or mouse anti-GFAP (1:500, Milipore MAB360), diluted in blocking solution. After washes in PBS, sections were incubated for 3 hours at room temperature with the corresponding secondary antibodies; goat anti-mouse AlexaFluor594 (1:300, Invitrogen A11032) and goat anti-rabbit AlexaFluor488 (1:300, Invitrogen A11034) for Figures 1 to 3 and donkey anti-rat AlexaFluor488 (1:500, A21208) and donkey anti-mouse AlexaFluor555 (1:500, A31570) for Figures 4 to 7. Nuclei were counterstained with DAPI (2 μg/mL in PBS) for 30 minutes. Sections were mounted onto glass slides and cover-slipped using FluorSave reagent (Millipore).

**For detection of other antigens (GFAP, Iba1, Tbr2),** an antigen retrieval step was performed prior to blocking by incubating the sections in 10 mM citrate buffer (pH 6.0) at 60°C for 30 minutes. The slices were then permeabilized in 1% PBST for 30 minutes at RT. Blocking buffer solution was prepared using 5% normal goat serum (Gibco, 1623 11530526) in PBST1% and incubated in the blocking buffer at RT for 1 hour. Slices were then incubated for 48 h at 4°C in PBST1% containing 5% of normal goat or donkey serum with the following primary antibodies: rabbit anti-GFAP (1:500, Dako Z0334), rat anti-Tbr2 (1:500, ebioscience, 14-4875-82) and goat anti-Iba1 (abcam ab5076). After 48 hours of incubation, the sections were rinsed 3 times for 10min in PBST1% and incubated for 3h in either of the following secondary antibodies in PBST1% and 5% of normal goat or donkey serum: donkey anti-rabbit AlexaFluor 647 (1:500, Invitrogen A31573), donkey anti-rat AlexaFluor 488 (1:500, Invitrogen A21208), donkey anti-goat AlexaFluor 555 (1:500, Invitrogen A21432) or goat anti-rabbit Alexa-594 (1:500, Life Technologies, A11037), goat anti-rabbit Alexa-488 (1:500, Invitrogen, A11034), and goat anti-rat Alexa Fluor 488 (1:500, Life Technologies, A11006). Sections were then rinsed 10min in PBS1x and incubated in 4,6 diamidino-2-phenylindole (DAPI, 1:1000) for 30 min to reveal nuclei before being rinsed again 3 times for 10min in PB 0.1M. The slides were dried completely and mounted using mounting medium (Sigma Aldrich, 345789) and sealed carefully using coverslips. Once dry, the slides were stored at 4°C until microscopy imaging.

**For Thioflavin-S** staining, sections were first processed for Iba1 immunostaining as described above, up to the mounting step. Sections were then mounted on gelatin-coated slides (Superfrost PLUS, Epredia, J1810AMNZ) and air-dried in the dark overnight. Slides were washed three times for 3 minutes in 50% ethanol, rinsed in sterile water for 3 minutes, and incubated in 1% Thioflavin-S solution (Sigma, T-1892) for 10 minutes. Excess dye was removed by sequential washes: five times for 3 minutes in 70% ethanol, three times for 3 minutes in 50% ethanol, and twice for 15 minutes in sterile Milli-Q water. Slides were air-dried for 30 minutes, coverslipped with antifade mounting medium (Sigma-Aldrich, 345789), and stored at 4°C in the dark until imaging. This protocol was adapted from published methods for amyloid detection using Thioflavin-S staining (Christensen and Pike, 2020).

#### Imaging

For cell quantification and imaging in Figures 1 and 2, images were acquired using an epifluorescence microscope (Zeiss AxioSkop2 Plus) equipped with a camera (Zeiss AxioCam MRm) and a 20× objective. All cells were manually quantified throughout the full thickness of each section. The dentate gyrus (DG) of each section was traced using ImageJ software to calculate DG volume. Sections immunolabeled for GFAP, Iba1, Nestin-GFP, Tbr2, or NeuN were imaged either with a laser-scanning confocal microscope (Zeiss LSM780 GaAsP, 20× objective, NA 1.0) using sequential channel acquisition or with a Nikon NI-E spinning disk confocal microscope (20× objective). For both systems, z-stacks were acquired through the full thickness of the tissue with adjacent optical sections (z-step) sufficient to resolve individual cells. Images from the Zeiss confocal were analyzed using Zen Blue software.

For cell quantification and imaging in Figures 3–5, all images were acquired on the Nikon NI-E spinning disk microscope with a 20× objective under identical acquisition settings across experimental conditions. Z-stacks were collected through the entire tissue volume, and images were analyzed using the NIS-Elements GA3 analysis module (Nikon Instruments, Melville, NY, USA). Custom GA3 pipelines were created to extract relevant morphological and fluorescence parameters. The Bright Spot function was used for cell quantification (BrdU, Iba1), and the Threshold function was applied for DCX^+^ immunolabeling to calculate volume fractions and for fluorescence intensity measurements (Iba1, GFAP). Threshold-based three-dimensional reconstructions of microglia around plaques were also performed to extract morphological features. Regions of interest (ROIs) were manually defined for each image to ensure analysis within appropriate anatomical areas (e.g., granule cell layer for BrdU, or granule cell layer, hilus, and molecular layer for Iba1 and GFAP). Once the custom analysis pipelines were established, images were automatically processed with experimental supervision to ensure accuracy.

**Figure 5.**
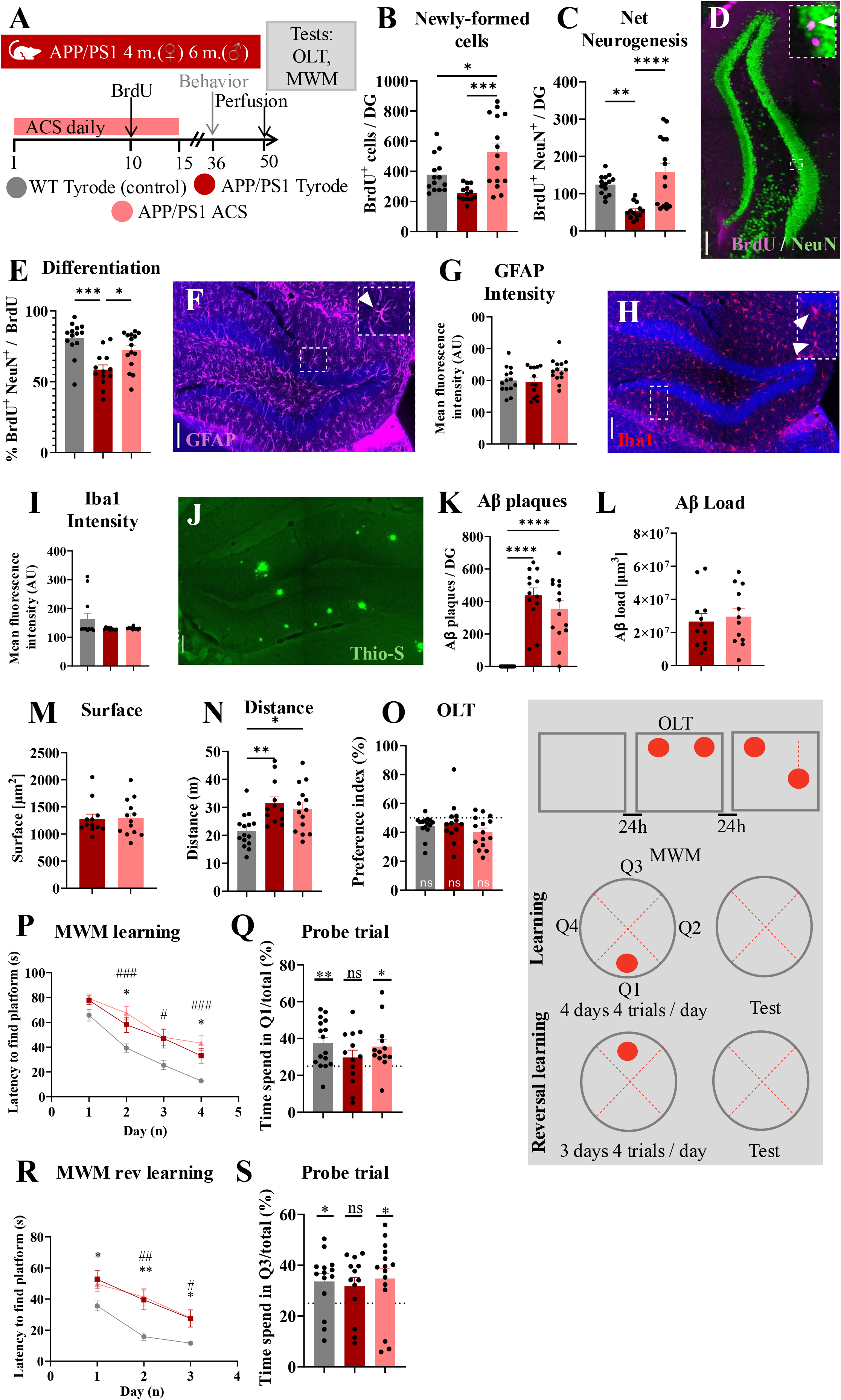
ACS rescues net neurogenesis and spatial memory without modifying amyloid load. **A)** Experimental timeline. **B)** Quantification of BrdU^+^ cells in the subgranular zone of the dentate gyrus (n = 42, F(2,39) = 10.39, p = 0.0002). **C)** Quantification of BrdU⁺/NeuN⁺ double-labelled cells in the dentate gyrus (n = 42, F(2,39) = 11.56, p = 0.0001). **D)** Representative confocal image. BrdU (Pink), NeuN (green). Inset show higher magnification of labelled cells with white arrow indicating an example of a BrdU-positive cell. **E)** Percentage of BrdU^+/^NeuN^+^ double-labelled cells from the total BrdU^+^ cell population (n = 42, F(2,39) = 10.95, p = 0.0002). **F)** Representative confocal image. GFAP (pink). Inset show higher magnification of labelled cells with white arrow indicating an example of a GFAP-positive cell. **G)** Mean fluorescence intensity of GFAP immunolabeling (n = 41, F(2,38) = 3.22, p = 0.05). **H)** Representative confocal image. Iba1 (red). Inset shows higher magnification of labelled cells with white arrows indicating an example of Iba1-positive cells. **I)** Mean fluorescence intensity of Iba1 immunolabeling (n = 34, F(2,30) = 2.43, p = 0.14). **J)** Representative confocal image. Thio-S (green). **K)** Quantification of amyloid-β plaques in the dentate gyrus (n = 41, F(2,38) = 33.94, p < 0.0001). **L)** Amyloid-β load in the dentate gyrus (n = 24, t = 0.4339, p = 0.67). **M)** Quantification of amyloid-β plaques Surface [µm^2^] (n = 25, t = 0.099, p = 0.92). **N)** Distance traveled (m) during the habituation phase of the open-field test (n = 41, F(2,38) = 6.79, p = 0.003). **O)** Preference index (%) in the Object Location Test (n = 43, F(2,40) = 1.32, p = 0.279). **P)** Escape latency (s) to reach the hidden platform across 5 days of acquisition training in the MWM (Day Factor (F(2.77, 110.8) = 66.7, p< 0.0001), Group factor (F(2,40) = 12.07, p < 0.0001) and Day x Group Factor (F(6, 120) = 1.03, p = 0.41)). **Q)** Percentage of time spent in the target quadrant during the probe trial (n = 42, F(2,39) = 1.25, p = 0.299). **R)** Escape latency (s) to reach the hidden platform across 3 days of acquisition in reverse learning in the MWM (Day Factor (F(1.3, 77.2) = 32.80, p< 0.0001), Group factor (F(2,40) = 8.95, p = 0.0006), and Day x Group Factor (F(4, 80) = 0.8, p = 0.80)). **S)** Percentage of time spent in the target quadrant during the reverse learning probe trial (n = 42, F(2,39) = 0.18, p = 0.835). Data was analyzed by two-way ANOVA (row factor × column factor) or one-way ANOVA, followed by Tukey’s multiple comparisons test. Significance is indicated as: * APP/PS1 Tyrode vs. WT Tyrode; # APP PS1 ACS vs. WT Tyrode. P-values: n.s. not significant, * p ≤ 0.05, ** p ≤ 0.01. Scale bar = 100 μm. OLT = Object Location Test. Morris Water Maze = MWM.

### Behavioral Tests

#### Memory Tests

##### T-Maze

The T-Maze test was used to assess spatial working memory and spontaneous alternation behavior. The apparatus consisted of a start arm (30 cm long) and two goal arms (30 cm long each) arranged in a T-shape, constructed from gray Plexiglas with walls 15 cm high and 10 cm wide, illuminated at 20 lux. During each trial, mice were placed at the start of the stem arm and allowed to choose freely between the left and right goal arms. After entering one arm, the mouse was confined for 30 seconds, then returned to the start arm for the next trial after a 10-second inter-trial interval. A testing session consisted of 10 consecutive trials. The percentage of spontaneous alternation (choosing the arm opposite to the previous choice) was calculated as an index of spatial working memory, with higher alternation rates indicating intact memory function. The apparatus was cleaned with 70% ethanol solution between subjects.

##### Y-Maze

The Y-maze test was used to assess spatial recognition memory based on the innate tendency of mice to explore novel environments. The apparatus consisted of three identical arms (43 × 14 × 23 cm) made of gray Plexiglas, illuminated at 20 lux. During the acquisition phase, mice were placed at the end of one arm with access to only two arms for 5 minutes, while the third arm remained blocked. After an 8-hour inter-trial interval, mice were returned to the maze with all three arms accessible for 3 minutes (retrieval phase). The preference index was calculated based on time spent in the novel arm versus familiar arms. Parameters recorded included first arm choice, time spent in each arm, number of entries per arm, and total arm entries as a measure of locomotor activity.

##### Object Location Test (OLT)

The OLT assessed spatial memory by evaluating the ability of mice to recognize changes in object location within a familiar environment. The test was conducted in a square arena (40 × 40 cm) over three consecutive days. A visual cue (black tape) was placed in the corner opposite to the objects to aid spatial orientation. On day 1 (habituation), mice explored the empty arena for 10 minutes. On day 2 (acquisition), two identical objects (A1 and A2) were placed in opposite corners, and mice explored for 10 minutes to establish familiarity. A discrimination index was calculated to identify any baseline object preference. On day 3 (test), copies of both objects were presented, with one object (A2, the non-preferred object to avoid bias) displaced 20 cm diagonally from its original position while the other remained stationary. Mice explored for 10 minutes, and the preference index was calculated as: (time exploring displaced object − time exploring stationary object) / total exploration time. The arena and objects were cleaned with 70% ethanol solution between trials to eliminate olfactory cues. Object positions and roles were counterbalanced across subjects.

##### Morris Water Maze (MWM)

Spatial learning and memory were assessed using the MWM in a circular pool (diameter: 140 cm) filled with water maintained at 24°C ± 1°C. The water was made opaque by adding 1 L of milk daily, rendering a circular platform (diameter: 12 cm) invisible 0.5 cm below the water surface. Four distinct visual cues were positioned around the pool to provide spatial references. Mice underwent acquisition training consisting of 4 trials per day for 4 consecutive days, with each trial lasting a maximum of 90 seconds. Mice were rescued immediately upon reaching the platform or, if they failed to locate it within 90 seconds, were gently guided to the platform.

Mice remained on the platform for 15 seconds to reinforce spatial learning. After each trial, mice were placed in a box with bedding under a warming light to dry and rest, with a minimum inter-trial interval of 25 minutes. Mice were released from a different quadrant in each trial, and the release order was randomized. On day 5, a probe trial was conducted with the platform removed to assess spatial memory retention. Subsequently, reversal learning was performed for 3 days with the platform relocated to the opposite quadrant, followed by a second probe trial on day 4. The pool was thoroughly cleaned after each daily session. Performance was tracked using video software (AniMaze), recording parameters including escape latency, swim speed, path length, and time spent in target quadrant during the probe trials.

### Multiplex analysis of mice serum

Cytokine concentrations in mouse serum were quantified using the Meso Scale Discovery (MSD) U-PLEX® platform (Meso Scale Diagnostics, Rockville, MD, USA). A custom 5-plex panel was employed to measure TNF-α, IL-1β, IL-10, IFN-γ, and MCP-1 according to the manufacturer’s instructions. Serum samples were diluted to ensure values fell within the linear dynamic range of the assay. Each plate included a standard curve and quality control samples. Data were acquired on an MSD instrument (e.g., MESO QuickPlex SQ 120, Meso Scale Diagnostics) and analyzed with the proprietary MSD Discovery Workbench® software (Meso Scale Diagnostics).

### Data processing, analysis and statistics

#### Behavioral Data

All behavioral parameters were acquired using an automated video tracking system (ANY-Maze, version 7.0, Stoelting Co., Wood Dale, IL, USA). The system automatically recorded and quantified behavioral parameters including distance traveled, time spent in defined zones, velocity, and entries into areas of interest. Critical parameters such as escape latency in the MWM were also measured manually to ensure accuracy and data validation. Raw data were exported from ANY-Maze as CSV files and processed using Python (Python Software Foundation, version 3.10) with the Pandas library (pandas-dev/pandas, https://pandas.pydata.org/) for organization by treatment group and statistical preparation. For the MWM, videos in which the tracking system failed to detect the animal during acquisition were re-analyzed post-hoc using manual scoring.

#### Imaging Data

Microscopy images were analyzed using the NIS-Elements GA3 analysis module (Nikon Instruments Inc., Melville, NY, USA). Custom analysis pipelines were created within GA3 to extract morphological and fluorescence parameters of interest. The resulting data were exported as CSV files and processed using Python (Python Software Foundation, version 3.10) with the Pandas library (pandas-dev/pandas, https://pandas.pydata.org/) to remove outliers (considered as larger or smaller than two times the standard deviation) and compute means per animal or experimental condition as appropriate.

#### Statistics

All statistical tests and graph generation were performed using GraphPad Prism (GraphPad Prism 9, GraphPad Software, San Diego, CA, USA). A critical probability of p < 0.05 was considered statistically significant. The specific statistical analyses performed for each experiment are indicated in the corresponding figure legends, including the type of test, sample size (’n’), and p-values. All values are presented as mean ± S.E.M.

## 3. Results

### 3.1. Astrocyte-conditioned solution enhances neurogenesis and improves memory in healthy conditions

We hypothesized that astrocytes secrete molecules that increase adult hippocampal neurogenesis and improve mood regulation and cognition. To test this possibility, murine primary hippocampal astrocytes were incubated in Tyrode solution for 24 hours for conditioning. The conditioned solution was then collected and filtered through a 0.22 µm filter (Millipore) to remove cellular debris, as previously described^53^ (**Figure 1a**). We then injected ACS or Tyrode (as control) solution i.p. daily for 8 days to C57/Bl6 8-week-old mice. One day after the last ACS injection, animals were injected with BrdU (100 mg / Kg), to enable the identification of dividing cells and were perfused one day later (**Figure 1b**).

Quantification of BrdU^+^ cells in the subgranular zone of the dentate gyrus (DG) revealed that ACS significantly increased cell proliferation as compared to vehicle (**Figure 1c, d**). ACS treatment did not modify Iba1^+^ microglia density, Iba1 fluorescence intensity (which integrates cell number and fluorescence intensity per cell) and GFAP fluorescence intensity (**Figure 1e–h**), suggesting that ACS did not affect inflammation.

To test the possibility that ACS affected the number of adult hippocampal stem cells, we used Nestin-GFP mice, in which neural stem cells express the green reporter protein GFP. Using the same ACS injection timeline (**Figure 1b**), we observed that ACS did not alter the number of Nestin^+^ stem cells. However, ACS significantly increased the number of Tbr2^+^ intermediate neural progenitor cells (**Figure 1i–j**).

Based on these findings, we next tested whether the expansion of Tbr2^+^ cells translated into increased net neurogenesis and behavioral changes. C57BL/6 mice were treated with ACS for 8 days, followed by BrdU labeling 1 day later. After a 3-week survival period—chosen to allow the maturation of newly generated neurons into their critical period of enhanced plasticity — we conducted a battery of behavioral tests of working memory (T-maze, Y-maze) and of pattern separation spatial memory (Object Location Test (OLT)^54^), from 28 to 35 days after the beginning of ACS injection. Mice were perfused on day 36 and prepared for histology (**Figure 1k**).

Histological analysis showed that ACS treatment significantly increased net neurogenesis, reflected by a higher number of BrdU^+^ NeuN^+^ cells (**Figure 1l, m**). We then assessed spatial working memory using the T-maze and Y-maze tests and found that ACS-treated and control mice displayed similar performances (**Figure 1n–o**). In the OLT, we did not find a significant difference between group. However, when the preference index was compared to chance level (i.e. 50%), we found that ACS-treated mice exhibited a preference for the displaced object (preference index of 62.94 ± 2.55 %, p = 0,0004, one-sample t-test) whereas Tyrode-treated mice did not (57.43 ± 3.64, p = 0,066) (**Figure 1p**).

Overall, ACS treatment promoted hippocampal neurogenesis and slightly improved spatial memory performance in a pattern-separation task.

### 3.2. Astrocyte-conditioned solution enhances neurogenesis and memory performance in aged mice

Physiological aging is accompanied with reduced AHN^10, 12^ and reduced cognitive performances^19^. Having established that ACS enhanced neurogenesis and memory in young mice, we next examined whether similar effects occur in aged mice. To this end, we administered ACS or Tyrode’s solution to 18-month-old C57BL/6 mice for 8 days. BrdU was injected on day 9 to label dividing cells, and brains were collected on day 10 (**Figure 2a**). Histological analysis revealed that ACS increased the number of proliferating cells in the subgranular zone of the DG (**Figure 2b, c**). Using the microglial marker Iba1, we found that ACS treatment did not alter either Iba1^+^ microglia density (**Figure 2d, e**) or the mean fluorescence intensity of Iba1^+^ labelling **(Figure 2f**).

To determine whether the effect of CS on cell proliferation translated into increased net neurogenesis and improved cognitive performance, we treated 18-month-old mice with ACS for 8 days, administered BrdU on day 9, and allowed a 28-day survival period to capture the integration of newly formed neurons during their window of plasticity^6^. Between days 28 and 33 following the first ACS injection, mice underwent a battery of cognitive tests (OLT, Y-maze) and were subsequently sacrificed on day 33 (**Figure 2g**). Histological assessment showed that ACS increased the total number of BrdU-labeled newly formed cells in the subgranular zone (**Figure 2h**). Consistently, we observed an increase in the number of immature DCX^+^ cells following ACS treatment (**Figure 2i, j**). Moreover, ACS treatment significantly increased net neurogenesis—quantified as BrdU^+^ cells co-expressing NeuN (**Figure 2k, l**), but not the neuronal differentiation of newly-formed cells (**Figure 2m**), indicating that the increased neurogenesis was due to an increase in cell proliferation but not in neuronal differentiation.

Given these pro-neurogenic effects, we evaluated whether ACS improved cognitive performance in aged mice in the OLT. ACS-treated mice identified the displaced object above chance (73.32 ± 8.122 %, p = 0.03) whereas controls did not identify the displaced object (48.82 ± 10.97 %, p = 0.92), despite no differences in the discrimination index between groups (**Figure 2n**). No significant difference between groups was detected in the Y-maze (**Figure 2o**), suggesting that ACS did not affect working memory.

Together, these results show that ACS enhanced neurogenesis in the dentate gyrus of aged mice and improved spatial memory performance.

### 3.3. Astrocyte-conditioned solution rescues adult neurogenesis and cognition in a mouse model of tau pathology

The pathological accumulation of phosphorylated tau within intracellular tangles is one of the major hallmarks of AD and several other neurodegenerative diseases. To determine whether ACS can enhance neurogenesis and ameliorate cognitive deficits in a tauopathy context, we used the THY-Tau22 mouse model, which expresses human 4R1N tau carrying the G272V and P301S mutations under the THY1.2 promoter^52^. This mouse models human Alzheimer’s-like tauopathy, including Tau hyperphosphorylation (AT8, AT100, AT180), and neurofibrillary tangle-like inclusions (Gallyas/MC1-positive). In addition, it exhibits non-spatial memory impairments by around 6 months of age, without motor deficits^52^. However, spatial memory deficits typically become evident only later, around 9 months of age^55^. Considering this, we treated 11–14-month-old THY-Tau22 mice with ACS or Tyrode solution for 8 consecutive days, and subsequently administered BrdU to label proliferating cells. In addition, aged-matched wild type (WT) littermates injected with Tyrode solution were used as controls. We then allowed a 24-day survival period to capture the integration of newly formed neurons during their window of plasticity and performed memory assessments (OLT and Y-maze) between days 24 and 32 after the first ACS injection. Mice were perfused-fixed on day 33 and prepared for histology (**Figure 3a**).

Histological quantification of BrdU^+^ cells in the DG showed a slight, but not significant, decrease in cell proliferation in THY-Tau22 mice as compared to WT or ACS-treated THY-Tau22 mice (**Figure 3b-d**). However, the number of BrdU^+^/NeuN^+^ double-labelled newborn neurons was decreased in THY-Tau22 mice and significantly increased by ACS treatment (**Figure 3e**), suggesting that ACS administration restored adult neurogenesis to a level comparable to WT mice.

To assess inflammation, we labelled microglial cells using Iba1 and quantified mean fluorescence intensity (**Figure 3f**). THY-Tau22 mice showed increased Iba1 immunoreactivity, which was mitigated by ACS treatment (**Figure 3g**).

When evaluating working memory performance in the Y-maze, we found that THY-Tau22 mice failed to significantly discriminate the closed arm (51.52 ± 5.21 %, p = 0.776) compared to their WT littermates (61.31 ± 4.23 %, p = 0.016), whereas this deficit was rescued by ACS treatment (68.61 ± 6.85 %, p = 0.035) (**Figure 3h**). Similarly, THY-Tau22 mice displayed an impairment at discrimination of displaced object in the OLT (51.91 ± 4.6 %, p = 0.687) as compared to WT littermates (62.18 ± 4.78 %, p = 0.019), and this discrimination index was restored by ACS, with treated animals exploring the displaced object significantly more than chance level (65.04 ± 4.94 %, p = 0.019) (**Figure 3i**).

Together, these results show that ACS treatment rescues both the neurogenesis impairment and the memory deficits associated with tau pathology in THY-Tau22 mice.

### 3.4. Astrocyte-conditioned solution rescues neurogenesis and reduces neuroinflammation in a mouse model of amyloidosis

Amyloid-β deposition has been proposed as an initiating event in disease progression^56^, ultimately contributing to neuronal loss and dementia. The APP/PS1 mouse model is a double-transgenic line carrying mutations in the APP and PS1 genes, resulting in amyloid-β plaque accumulation in the hippocampus starting at approximately 4 months of age ^57, 58^. This model of cerebral amyloidosis has been reported to exhibit impairments in AHN as early as 2 months of age^36^.

To investigate the effect of ACS treatment on AHN impairments in APP/PS1 mice, we used female APP/PS1 mice at 4 months of age and male APP/PS1 mice at 6 months of age, that display comparable amyloid-β loads^59^, as well as their age-matched and sex-matched WT littermates. Mice received intraperitoneal injections of ACS or Tyrode’s solution for 15 consecutive days and were assigned to three experimental groups: WT mice treated with Tyrode’s solution (WT Tyrode), APP/PS1 mice treated with Tyrode’s solution (APP/PS1 Tyrode), and APP/PS1 mice treated with ACS (APP/PS1 ACS). To label proliferating cells, BrdU was administered on day 10 of the treatment period, and animals were perfused-fixed on day 15 (**Figure 4a**).

We first examined whether ACS differentially affects neurogenesis and astrogenesis, as astrogenesis is known to be altered in APP/PS1 mice^60^. Quantification of BrdU-labeled cells revealed a significant reduction in cell proliferation and early cell survival in the DG of APP/PS1 mice compared with WT controls. Importantly, this reduction was rescued by ACS treatment (**Figure 4b**). Analysis of dentate gyrus volume revealed no differences among groups (data not shown), indicating that changes in progenitor proliferation occurred independently of DG volume.

In addition, APP/PS1 mice exhibited an increased proportion of BrdU⁺ cells expressing GFAP and displaying a stellate morphology compared with WT mice, consistent with previous reports of enhanced astrogenesis. This increase in astrogenic fate was normalized to WT levels following ACS treatment (**Figure 4c, d**). We next assessed the impact of ACS on immature neurons by quantifying the DCX⁺ volume fraction, defined as the proportion of the dentate gyrus occupied by DCX⁺ immunolabeling (**Figure 4e–j**), which integrates both cell number and dendritic length. APP/PS1 mice showed a marked reduction in DCX⁺ volume fraction, which was fully restored by ACS treatment (**Figure 4k**).

To determine whether ACS affected markers of neuroinflammation in the dentate gyrus of APP/PS1 mice, we quantified the number and fluorescence intensity of GFAP⁺ and Iba1⁺ cells, markers of astrocytes and microglia, respectively. Quantification of GFAP⁺ cells revealed an increased number of astrocytes in the dentate gyrus of APP/PS1 mice compared with WT littermates. This increase was no longer statistically significant in ACS-treated APP/PS1 mice (**Figure 4l,m**). Consistently, GFAP fluorescence intensity was elevated in APP/PS1 mice regardless of treatment (**Figure 4n**).

Similarly, the number of Iba1⁺ cells was increased in APP/PS1 mice compared with WT littermates, whereas no significant increase was observed in ACS-treated APP/PS1 mice (**Figure 4o,p**). In contrast, Iba1 fluorescence intensity was elevated in APP/PS1 mice irrespective of treatment (**Figure 4q**). Together, these data indicate increased GFAP and Iba1 immunoreactivity in the dentate gyrus of APP/PS1 mice, with a trend toward attenuation following ACS treatment.

Analysis of amyloid-β plaque with Thioflavin-S labeling confirmed that APP/PS1 mice exhibited plaques in the DG (**Figure 4r,s**). However, ACS treatment did not reduce plaque number, plaque surface, or overall plaque load (**Figure 4s,t,u**).

Previous data from our laboratory indicate that ACS has anti-inflammatory properties^44^. To further assess inflammation in these experimental groups, we quantified circulating cytokine levels. Plasma IL-1β concentrations showed a trend toward higher levels in APP/PS1 mice and lower levels following ACS treatment, approaching those observed in WT mice (**Figure 4v**). A similar pattern was observed for TNF-α and MCP-1, both of which were increased in APP/PS1 mice and reduced by ACS treatment (**Figure 4w,x**). Although none of these differences reached significance level, their common trend points to increased inflammation in APP-PS1 mice and reduced inflammation with ACS treatment. In contrast, no differences were detected between groups for IFN-γ or IL-10 levels (**Figure 4y,z**).

Collectively, these results indicate that ACS restores cell proliferation, astrogenesis and DCX⁺ cells number and/or morphology in the DG of APP/PS1 mice and induces a trend towards a reduction in circulating pro-inflammatory markers.

### 3.5. Astrocyte-conditioned solution reduces activation of plaque-associated microglia

Microglia are the resident immune cells of the brain and play a complex role in Alzheimer’s disease progression. Recent studies have identified a distinct microglial subtype, termed disease-or plaque-associated microglia, characterized by altered transcriptional profiles and implicated in disease progression^61^. Since ACS showed anti-inflammatory effects (**Fig. 4v-z**), we investigated whether ACS treatment may modulate the morphological state of plaque-associated microglia.

To this aim, we used the same cohort of animals described in Figure 4a, which received ACS treatment for 15 days prior to sacrifice (**Supplementary Figure 1a**). Microglial morphology was assessed using three-dimensional reconstructions derived from confocal z-stack images (**Supplementary Figure 1b–d**), and multiple morphological parameters were quantified.

Analysis of non–plaque-associated microglia—defined as cells not in direct contact with amyloid-β plaques—revealed no significant differences in diameter, perimeter, length, surface area, circularity, elongation, or shape factor among WT, APP/PS1 Tyrode, and APP/PS1 ACS groups (**Supplementary Figure 1e–k**).

We next analyzed microglia in direct contact with amyloid-β plaques (**Supplementary Figure 1l**) using three-dimensional reconstructions (**Supplementary Figure 1m,n**). A comparison of plaque-associated microglia from ACS-and Tyrode-treated APP/PS1 mice revealed that ACS treatment significantly altered microglial morphology. Specifically, plaque-associated microglia from ACS-treated mice exhibited reduced perimeter, area, diameter, and process length compared with Tyrode-treated controls, consistent with a less activated morphological phenotype (**Supplementary Figure 1o–u**).

Collectively, these findings indicate that ACS selectively modulates plaque-associated microglia, attenuating morphological features associated with a highly activated microglial state.

### 3.6. Astrocyte-conditioned solution rescues net neurogenesis and spatial memory without modifying amyloid load

To determine whether the effects of ACS on cell proliferation translated into long-lasting changes in net neurogenesis and cognitive performance, we treated an independent cohort of APP/PS1 mice and WT littermates with intraperitoneal ACS injections for 15 days. BrdU was administered on day 10 to label proliferating cells. Following ACS treatment, a 3-week waiting period was included to coincide with the heightened plasticity window of newly generated neurons. Cognitive performance was then assessed between days 36 and 49 after the first ACS injection using the object location test (OLT) and the Morris water maze (MWM), and animals were perfused-fixed on day 50 (**Figure 5a**).

APP/PS1 mice exhibited a marked reduction in the number of newly formed, BrdU⁺ cells as compared to WT mice, which was rescued by ACS treatment (**Figure 5b**). Consistent with this, the number of newly-formed BrdU⁺/NeuN⁺ neurons was reduced in APP-PS1 mice (53.54 ± 5.9 cells/DG) as compared to WT mice (124.3 ± 7.01 cells/DG), which was rescued by ACS treatment (158 ± 23.62 cells/DG; **Figure 5c,d**). Analysis of neuronal differentiation, expressed as the percentage of BrdU⁺/NeuN⁺ cells relative to total BrdU⁺ cells, revealed reduced neuronal differentiation in APP/PS1 mice, which was restored to WT levels following ACS treatment (**Figure 5e**).

We next assessed neuroinflammatory markers in the dentate gyrus of this cohort. Quantification of GFAP and Iba1 fluorescence intensity revealed no significant differences between groups (**Figure 5f–i**), suggesting no persistent neuroinflammatory changes at this later time point. Amyloid-β plaque load in the dentate gyrus was assessed using Thioflavin-S labeling (**Figure 5j**). Consistent with our previous observations (**Figure 4r-u**), no significant differences in plaque number were observed between ACS-and Tyrode-treated APP/PS1 mice, although a trend toward reduced plaque number was apparent in ACS-treated animals (353.7 ± 52.19 vs. 437.7 ± 46.18 plaques; **Figure 5k**). Amyloid plaque load and plaque surface were also unchanged by ACS treatment (**Figure 5l,m**).

Finally, we evaluated behavior and cognitive performances. In the open-field test, APP/PS1 mice exhibited increased locomotor activity as compared to WT mice, which was not reduced by ACS treatment (**Figure 5n**). In the OLT, the discrimination index was similar between groups and at chance level, suggesting that all animals failed at discriminating the displaced object in these conditions (**Figure 5o**).

In the MWM, all experimental groups displayed significant learning across training days, reflected by progressive reductions in escape latency (two-way repeated-measures ANOVA, main effect of day, p < 0.001). APP/PS1 mice exhibited overall longer escape latencies compared to WT mice, regardless of treatment (main effect of genotype, p < 0.001; **Figure 5p**). To test for spatial memory, we performed a probe trial 24 hours after the last training. WT mice showed a significant preference for the target quadrant, whereas Tyrode-treated APP/PS1 mice performed at chance level. Notably, ACS-treated APP/PS1 mice exhibited a restored preference for the target quadrant, indicating improved memory retention (**Figure 5q**). To test for cognitive flexibility, we then performed a reversal learning test. All groups showed significant learning across training days (main effect of day, p < 0.001; **Figure 5r**), although APP/PS1 mice continued to display increased escape latencies regardless of treatment (main effect of genotype, p < 0.001). In the reversal probe trial (**Figure 5s**), APP/PS1 mice failed to show a significant preference for the target quadrant, unlike WT mice. Importantly, this deficit was rescued by ACS treatment, as ACS-treated APP/PS1 mice spent significantly more time in the target quadrant.

Together, these findings demonstrate that ACS rescues hippocampal neurogenesis deficits and improves spatial memory performance in APP/PS1 mice without altering amyloid burden.

## 4. Discussion

In the present study we tested the effect of the astrocytic secretome, in the form of ACS, on the hippocampal neurogenic niche, adult neurogenesis and memory performances. First, we found that ACS administration in young healthy mice increased the Tbr2⁺ progenitor cell population, resulting in an overall increase in cell proliferation and net neurogenesis and a mild improvement in memory performance. Next, we found that ACS treatment also increased hippocampal cell proliferation and net neurogenesis in aged mice, a context in which AHN typically declines, and significantly improved memory performance in the OLT. In both young and aged healthy mice, ACS did not induce detectable changes in inflammatory markers. We next examined the effects of ACS in two distinct mouse models of Alzheimer’s disease (AD). In the THY-Tau22 tauopathy mouse model, ACS treatment rescued net neurogenesis and restored memory performance in the OLT. THY-Tau22 mice exhibited increased Iba1 immunostaining in the dentate gyrus, which was mitigated by ACS administration. In the APP/PS1 model of amyloidosis, ACS rescued deficits in proliferation, net neurogenesis, and DCX⁺ volume fraction, confirming a robust pro-neurogenic effect. In this model, ACS also normalized increased astrogenesis. Although ACS did not improve performance in the OLT in APP/PS1 mice, it enhanced cognitive function in the MWM, rescuing both spatial memory and cognitive flexibility deficits relative to WT littermates. Notably, ACS treatment did not alter amyloid-β load, plaque number, or general inflammatory markers in the dentate gyrus; however, it reduced proinflammatory cytokine levels and decreased microglial activation in plaque-associated regions. Together, these results suggest that ACS restores neurogenesis and cognition in ageing and AD, independently from amyloid pathology.

A key aspect of this study is the origin of the ACS, which was derived from postnatal astrocytes. One plausible explanation for its robust pro-neurogenic effects is that the postnatal astrocyte secretome introduces youthful factors capable of rejuvenating the neurogenic niche ^62^. Indeed, postnatal astrocytes express higher levels of neurotrophic and pro-neurogenic molecules, including FGF2, D-serine, and other growth factors, which are reduced in adult or reactive astrocytes^63, 64^, indicative of diminished trophic support. We therefore propose that the age-associated decline in astrocyte-mediated support within the neurogenic niche can be partially compensated by administering homeostatic factors secreted by postnatal astrocytes. In support of this possibility, ACS induced a stronger effect on cell proliferation in young mice as compared to old mice (77,1% versus 51,13% increase, respectively). However, owing to the technical difficulty of maintaining aged astrocytes *in vitro*, we have not compared the role of aged ACS on the same mechanisms. It is therefore unclear whether the secretome from aged astrocytes may retain a similar function. Nevertheless, because healthy young adult mice are presumed to have functionally intact astrocytes, the stimulation of adult neurogenesis by postnatal ACS suggests that ACS may additionally act by supplementing pro-neurogenic molecules that are likely already present within the niche.

Interestingly, ACS produced a substantially larger increase in net neurogenesis in the two AD models examined, (neurogenesis increased by 182.78% in THY-Tau22 mice and 185.11% in APP/PS1 mice), as compared to wild-type animals (61.15% increase in young adult mice and 57.53% in aged mice). This pronounced difference of effect suggests that neurodegenerative conditions involve a more severe disruption of astrocytic function and trophic support, frequently accompanied by persistent neuroinflammation. Notably, elevated neuroinflammatory markers were observed in the dentate gyrus of THY-Tau22 and APP/PS1 mice but were absent in young and aged wild-type mice. Under these conditions, supplementation with homeostatic astrocyte-derived factors may more effectively compensate for these deficits, eliciting a markedly stronger pro-neurogenic response than in wild-type animals, where the effect likely reflects the intrinsic neurogenic capacity of the astrocyte secretome. Consistent with this interpretation, transcriptomic studies in AD have identified disease-associated astrocyte populations with reduced expression of neuronal support genes and increased activation of inflammatory pathways ^48, 49^. In addition, metabolic dysfunction in astrocytes during neurodegeneration may impair the release of gliotransmitters and trophic factors essential for neuronal and circuit homeostasis^65^. Together, these findings support the notion that supplementation with homeostatic astrocyte-secreted factors is particularly effective in neurodegenerative contexts and may also explain why ACS more robustly rescues cognitive performance in AD models, while its effects on memory and pattern separation in wild-type animals remain comparatively modest.

We recently showed that ACS produces a direct proneurogenic effect mediated by the release of molecules that target adult hippocampal stem/progenitor cells^53^. In addition, ACS induces anti-inflammatory effects that can also participate to its proneurogenic effect^44^. In the present study, microglia morphology, or its expression of Iba1 was not altered in aged animals, nor in mouse models of Alzheimer’s disease. In APP/PS1 mice however, plaque-associated microglia displayed strong morphological modifications. Indeed, the plaque microenvironment represents a particularly inflammatory niche. Amyloid-β plaques activate the microglial inflammasome and facilitate further deposition^66–68^. Within this context, a specialized microglial subtype— disease-associated microglia (DAM)—emerges, characterized by typical microglial markers but downregulated homeostatic genes^61, 69^. These DAMs co-localize with plaques^70^ and exhibit enhanced phagocytic capacity, upregulating lysosomal degradation, phagocytosis, and lipid metabolism^71^. Interestingly, evidence suggests a dual microglial role: initially seeding amyloid-β deposition, but subsequently compacting plaques and reducing dystrophic neurites at later stages^72^. We observed that ACS reduced the morphological modifications of plaque-associated microglia. In addition to its effect on microglia morphology, ACS mitigated the increase in several circulating inflammatory cytokines. Although this effect did not reach significance, the convergence of these effects with the morphological modifications of plaque-associated microglia suggests that ACS may also produce an anti-inflammatory effect in APP/PS1 mice both in the periphery and in the brain. However, whether the anti-inflammatory of ACS is beneficial for memory and neurogenesis in APP/PS1 mice remains to be demonstrated.

Here, we showed that ACS enhances cognitive performance across models of healthy, ageing, and AD mice, with the most consistent effects observed in tasks requiring pattern separation. While adult hippocampal neurogenesis has been closely linked to spatial processing and pattern separation^5^, ACS may also modulate cognition through neurogenesis-independent mechanisms, including effects on synaptic plasticity, astrocyte–neuron metabolic coupling, and inflammation-driven circuit dysfunction. Thus, the relative contribution of neurogenesis to the cognitive benefits of ACS cannot be fully resolved in the present study. Future studies selectively inhibiting neurogenesis will be necessary to determine its causality in the cognitive benefits of ACS.

Our findings show that ACS treatment improved cognitive performance in the Morris water maze in APP/PS1 mice, despite having no detectable effect on amyloid-β load, plaque number, or plaque morphology. New approved therapies for AD are largely based on targeting and removing amyloid-β^73^; however, the associated cognitive benefits remain modest ^74, 75^. This apparent disconnect between substantial amyloid reduction and limited cognitive improvement supports the need for combining therapeutic strategies. Specifically, our data argue in favor of approaches that combine amyloid-β–targeting antibodies with interventions capable of compensating for cellular dysfunction and restoring brain homeostasis, particularly those that enhance hippocampal plasticity and modulate neuroinflammation. By leveraging the intrinsic capacity of astrocytes to orchestrate neurogenesis and neurogenic niche homeostasis, ACS acts as a non-genetic, brain-derived, and niche-targeted modulator of plasticity across the lifespan. The particularly strong effects observed in neurodegenerative models highlight the vulnerability of astrocytic support mechanisms in disease and their potential for therapeutic rescue. Together, our findings identify the astrocytic secretome as a promising avenue to restore brain homeostasis and cognitive function in ageing and neurodegeneration.

## 6. Acknowledgements

The authors wish to thank Fulvio Magara, Fanny Thevenaz and Clara Rossetti for their help in the preparation and execution of the Morris Water Maze and Prof. Johannes Graeff from the Brain and Mind Institute at the EPFL for the generous donation of APP/PS1 mice.

## 9. Author contributions

MV, NT, FC, KR, EG: Conceptualization. MV, CC, FC, KV, KG, JN, EG: Data curation, Formal Analysis, Investigation and Software. MV, NT: Manuscript writing. NT: Supervision and Funding acquisition.

---

**Supplementary Figure 1.**
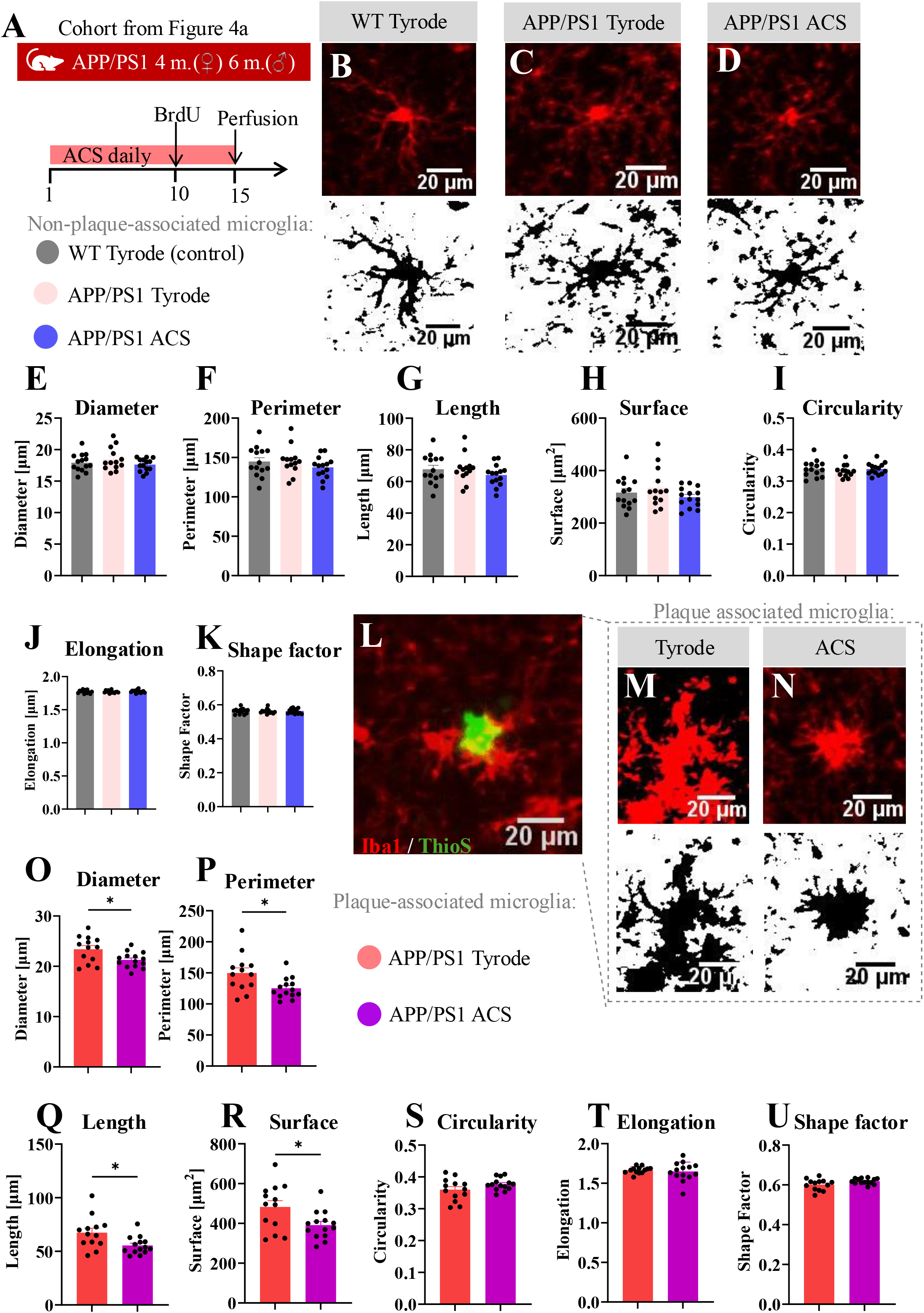
ACS reduces activation of plaque-associated microglia. **A)** Experimental timeline. Animals from the Figure 4a cohort. **B-D)** Representative confocal images (Iba1 (red)) and their corresponding threshold-processed images used for microglia morphology analysis (black). **E)** Microglia diameter [µm] (n = 41, F(2,38) = 1.2, p = 0.212). **F)** Microglia perimeter [µm] (n = 41, F(2,38) = 0.96, p = 0.383). **G)** Microglia length [µm] (n = 41, F(2,38) = 0.95, p = 0.395). **H)** Microglia surface [µm^2^] (n = 41, F(2,38) = 1.26, p = 0.295). **I)** Microglia circularity (n = 41, F(2,38) = 0.518, p = 0.599). **J)** Microglia elongation [µm] (n = 41, F(2,38) = 0.295, p = 0.746). **K)** Microglia shape factor (n = 41, F(2,38) = 0.014, p = 0.985). **L)** Representative confocal image. Iba1 (red), Thioflavin-S (green). **M-N)** Representative confocal images in plaque associated microglia (Iba1 (red)) and their corresponding threshold-processed images used for microglia morphology analysis (black). **O)** Microglia diameter [µm] (n = 27, t = 2.6, p = 0.016). **P)** Microglia perimeter [µm] (n = 27, t = 2.64, p = 0.014). **Q)** Microglia length [µm] (n = 27, t = 2.62, p = 0.015). **R)** Microglia surface [µm^2^] (n = 27, t = 2.49, p = 0.0198). **S)** Microglia circularity (n = 27, t = 1.58, p = 0.13). **T)** Microglia elongation [µm] (n = 27, t = 0.51, p = 0.615). **U)** Microglia shape factor (n = 27, t = 2.03, p = 0.053). Data is shown as mean ± SEM. Statistics: One-way ANOVA with Tukey’s multiple comparisons test or unpaired T-test. *p ≤ 0.05. Scale bar = 20 µm.

